# Evolutionary discovery and characterization of fungal transcriptional activators using active learning

**DOI:** 10.1101/2025.09.12.675635

**Authors:** Lucas Waldburger, Hunter Nisonoff, Marissa Zintel, Liam D. Kirkpatrick, Angelica Lam, Nathan Lanclos, Jay D. Keasling, Max V. Staller, Patrick M. Shih

**Author notes:** Corresponding authors. M.V.S. ; P.M.S.

## Abstract

Biological discovery and design are increasingly being guided by predictive models in place of costly experimentation. However, existing datasets are often biased by overrepresentation from model organisms, leading to failures in evolutionary studies of non-model species. We present a hybrid framework that leverages high-throughput molecular assays and active learning to quantify biological properties across evolutionary space. We focus on transcriptional activators, which contain activation domains (ADs) that promote gene expression. ADs are intrinsically disordered and poorly conserved, which limits their study using comparative genomics. Here, we developed ADhunter, a high-capacity regression model that outperforms state-of-theart algorithms in identifying and quantifying the strength of transcriptional activators. Model uncertainty was used to guide evolutionary sampling across 7.8 million proteins from 2,400 fungal genomes. We functionally characterized 9,836 ADs from 1,071 fungal genomes, providing a 15.5-fold expansion in genome representation compared to existing datasets. Comprehensive sampling from non-model genomes improved model generalizability and provides the first functional annotation for 3,416 proteins from 670 non-model fungi. Model interpretability analysis aligns with the biophysical model of AD function and reveals novel, underrepresented protein codes, highlighting the importance of sampling from non-model organisms to build evolutionarily robust models for predicting biological properties.

## Introduction

Advances in DNA synthesis and sequencing have enabled high-throughput molecular technologies that are transforming biological research [1, 2]. These methodologies have shifted experimentation from small-scale characterization of a few hypotheses to large-scale assays capable of evaluating hundreds or thousands of hypotheses in parallel. While small-scale experiments have elucidated gene-level principles, large-scale approaches enable researchers to uncover genome-level features governing biological functions [3, 4].

In biological discovery and design, high-capacity regression models are increasingly replacing costly and time-consuming functional characterization with cheap and fast inference [5–8]. These surrogate models approximate the behavior of a more complex or computationally expensive function, enabling estimation or inference by serving as a proxy for biological properties. High-throughput assays provide functional measurements for training surrogate models, but existing datasets are often biased by overrepresentation of sequences from model organisms, limiting model generalizability in evolutionary studies (Fig. 1A). Decades of sequencing and experimentation have highlighted that model organisms represent only a fraction of biological diversity. While experimental scientists rely on model systems to elucidate biological principles through standardized procedures, high-throughput assays are limited by the constraints of DNA synthesis not by organismal origin. Evolutionary space represents a broad distribution of functionally-enriched sequences, especially from non-model organisms, that challenge high-throughput technologies with edge cases, enabling a more rigorous evaluation of the assay’s capacity to quantify biological properties (Fig. 1B). Furthermore, most algorithms are known to suffer from pathologies such as overconfident predictions and reduced accuracy when performing inference on regimes far from the training distribution [9, 10] (Fig. 1C). Therefore, models trained exclusively on sequences from model organisms are unreliable when performing inference on divergent sequences from non-model species. Successful design of predictive models for evolutionary studies thus requires comprehensive training datasets to capture underrepresented relationships over a property of interest.

**Figure 1.**
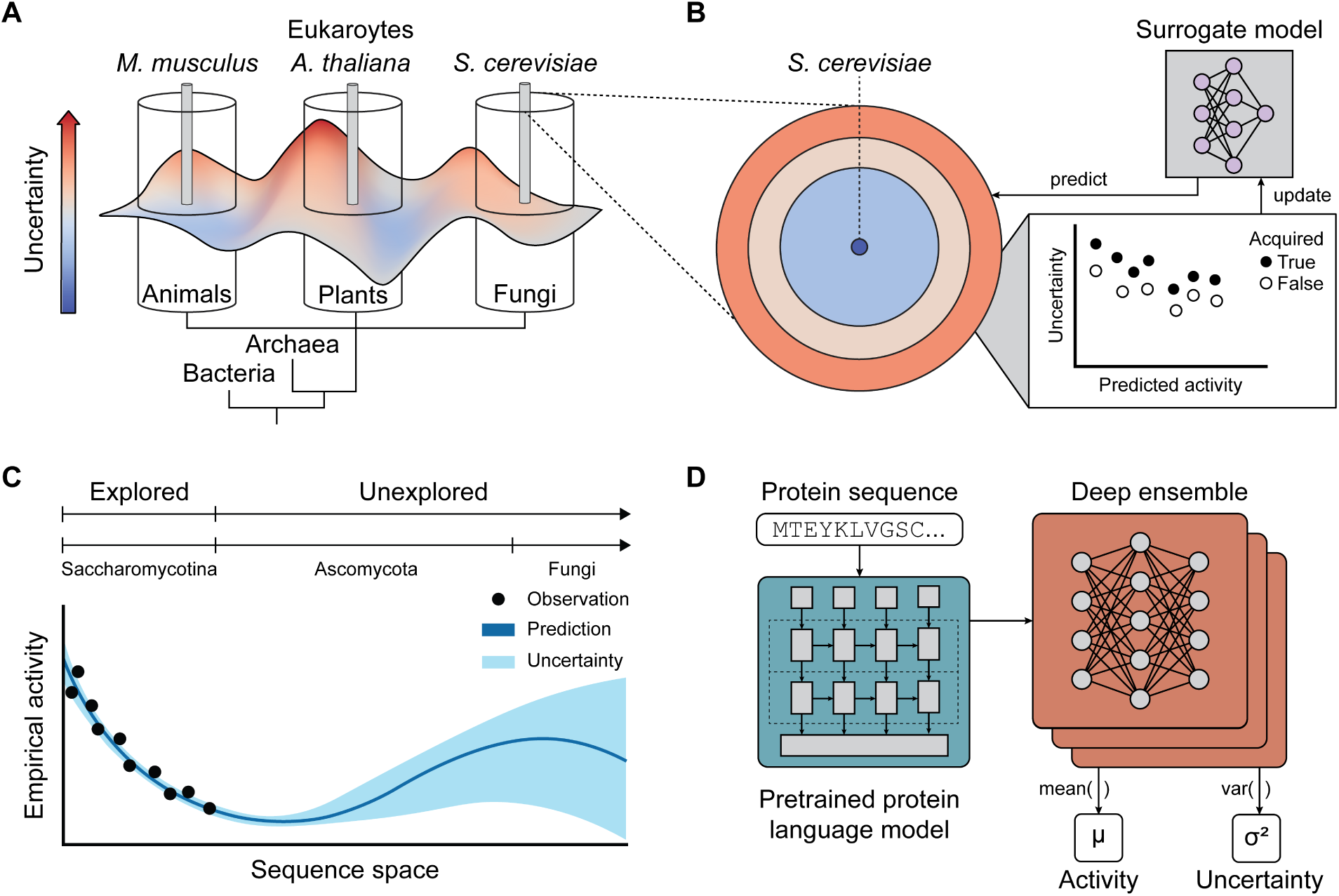
Quantitative sequence-to-function modeling of transcriptional activators. (A) Model organisms (e.g., *S. cerevisiae*, *A. thaliana*, *M. musculus*) are useful experimental systems, however, their sequence space represents a fraction of biological diversity for their respective branches of life. (B) Predictive models used to predict biological properties trained exclusively on sequences from model organisms perform poorly in evolutionary studies. Predictive models can be made into generalizable predictors of biological properties by acquiring labeled data with high uncertainty across evolutionary space and updating the initial model. (C) Underexplored regimes in sequence space have high uncertainty, thereby enabling experimentalists to prioritize functional characterization efforts to construct comprehensive training datasets. (D) ADhunter leverages neural encodings from pretrained protein language models to provide general, task-agnostic inputs and a deep ensemble to quantify AD activity as the mean predicted activity across individual models, and measure epistemic uncertainty based on the variance in predictions.

One such property is sequence-to-function prediction of transcriptional activators, which promote gene expression [11]. Activation domains (ADs) undergo dynamic interactions with components of transcriptional machinery to enhance expression of target genes. Predicting the extent to which ADs promote gene expression remains difficult due to their intrinsic disorder, multiple modes of binding, and poor sequence conservation, limiting comparative genomics approaches [12]. Small-scale characterization has failed to capture the mechanistic basis of many facets of transcriptional activation. Recently developed high-throughput technologies have enabled quantification of thousands of ADs in parallel [11, 13], providing functional measurements for training surrogate models. While existing models can identify key properties of AD sequences, most datasets are biased toward the model fungus, *Saccharomyces cerevisiae* [14, 15], and the model plant, *Arabidopsis thaliana* [16, 17]. Models trained on these datasets can identify a functionally-conserved class of ADs known as acidic ADs [18–20], but fail to detect less-characterized AD classes, limiting their generalizability in non-model genomes. Since ADs lack structural constraints, they can explore a much larger sequence space compared to structured proteins [21], making model generalizability imperative for evolutionary studies of gene regulation.

In this study, we leverage the growing volume of sequencing data from non-model genomes and active learning to discover transcriptional activators across fungal evolutionary space. Fungi are largely understudied, yet biosynthesize natural products that have transformed modern medicine [22]. Non-model fungi are increasingly appreciated for their ecological role in mediating plant health and decomposing organic compounds [23], which are being engineered for sustainability applications in bioremediation and food production [24]. Functional characterization has primarily focused on sequences from *Saccharomycotina*, while the broader protein diversity across *Ascomycota* and other fungal divisions remains largely uncharted. We developed ADhunter, a high-capacity regression model that outperforms the state-of-the-art model, TADA [25], at identifying and quantifying the strength of transcriptional activators (Fig. 1D). Machine-based uncertainty was used to predict activity for 7,842,516 proteins across 2,400 fungal genomes. The predicted activity and associated uncertainty guided the acquisition and downstream functional characterization of 9,836 proteins from 1,071 fungal genomes. We demonstrate how performing active learning on non-model genomes significantly enhanced ADhunter’s ability to quantify the activity of diverse transcriptional activators, especially non-acidic sequences that are leucine- and phenylalanine-enriched. Our results demonstrate how integrating high-throughput molecular technologies with active learning from non-model genomes enables scaling of genome-level characterization towards evolutionary-level functional genomics.

## Results

### Sequence-to-function modeling for precise quantification of transcriptional activators

Algorithms that are robust across evolutionary space require comprehensive training datasets to predict a biological property of interest. We sought to create a regression model to quantitatively predict AD activity. A quantitative model would enable accurate identification of AD boundaries and peak activity in natural sequences to study intrinsically disordered protein evolution [20], such as in non-model organisms that have evolved genetic regulation to biosynthesize natural products and adapt to ecological niches. Furthermore, a quantitative model can be used in protein engineering contexts to design transcription factors (TFs) for fine-tuned gene expression in synthetic biology. We used an initial dataset of 17,609 53 AA tiles from fungal and plant proteins that were previously characterized for AD activity using a high-throughput assay [16]. The GFP:mCherry ratio for 500,000 sorted cells was selected as the measurement of AD activity, based on the distribution of empirical activity (Fig. S1).

Existing AD predictors fail to accurately identify domain boundaries and quantify domain strength in activating gene expression [16]. While preparing this study, a new model, TADA [17], achieves state-of-the-art performance and has been shown to outperform first-generation models ADpred [14] and PADDLE [15]. However, their training performance is optimized using a classification objective, whereas transcriptional activation is continuous and should be modeled with a regression objective. We systematically evaluated protein sequence representations to enhance sample efficiency, using performance on a held-out test dataset as the benchmark. Functional characterization of AD sequences is costly and time-consuming, making it essential for predictive models to extract the most information from each datapoint. Binary (i.e., one-hot) and integer encodings of protein sequences are frequently used due to their simplicity; however, this approach fails to capture secondary structure and biochemical properties of amino acids [26, 27]. Alternatively, features from human-selected sequence descriptors may introduce biases. Combining both approaches can lead to unequal weighting and increased complexity, resulting in reduced performance (Table S1). For instance, PADDLE [15] performs well at identifying acidic ADs, but has poor quantitative performance on the initial dataset (Pearson r = 0.261; RMSE = 0.336). Excluding secondary structure predictions improves performance (Pearson r = 0.338; RMSE = 0.329). Continuous neural encodings from pretrained protein language models provide general, task-agnostic features that capture evolutionary signals [28–31]. We trained a convolutional neural network (CNN) and found that neural encodings from an evolutionary-scale protein language model (ESM) [32] outperformed one-hot and all other representations on a held-out test dataset (Pearson r = 0.744; RMSE = 0.664; Table S2). Since ADs are disordered and poorly conserved, we assessed whether neural encodings from ESM capture meaningful sequence features. We trained models using either the first or last layer embeddings from ESM on a held-out test set (Fig. S2). For ESM version 1, models using last-layer embeddings outperformed those using first-layer embeddings (p-value ≤ 0.0112, Student’s t-test with Bonferroni multiple comparison correction), suggesting that deeper representations better capture features relevant to AD function. In contrast, ESM version 2 models showed comparable performance across both layers, with higher overall accuracy. These results indicate that neural encodings from pretrained models outperform simple encodings and that pretrained protein language models learn representations that are informative for modeling disordered and poorly conserved regions like ADs.

After determining the protein sequence representation, we evaluated model architectures to ensure optimal performance of the AD prediction task. Neural encodings combined with lightweight regressors have been shown to outperform complex architectures trained on simpler features [33]. Therefore, we selected our model architecture based on the ability to balance simplicity and interpretability while maximizing performance on the held-out test dataset. We evaluated 44 lightweight regressors, all of which performed poorly on the held-out test dataset (Fig. S3). Several lightweight classifiers performed well indicating that the binary classification task is less challenging than the regression task (Fig. S4). More complex models incorporating a convolutional layer performed the best. In particular, a CNN with residual connections (ResNet) had the highest performance on the held-out test dataset (Fig. S5). The ResNet model with ESM encodings was named ADhunter. When trained on the initial dataset, ADhunter achieved better quantitative performance than TADA (Pearson r = 0.538; RMSE = 0.995; Table S3). An ablation study of the model architecture used in TADA reveals that the CNN layer alone is sufficient to achieve optimal predictive performance (Table S4). These results demonstrate how a ResNet trained with neural encodings can outperform a more complex architecture trained on human-selected features at quantitatively predicting AD activity.

### Deep ensembling enables uncertainty measurement and improves model generalizability

Quantifying uncertainty enables researchers to distinguish between confident predictions and those that are unreliable, guiding prioritization of candidates for experimental validation and exploration of uncharacterized sequence space. Pathological failure becomes especially problematic when performing inference on sequences that are far from the training distribution [33, 34]. Existing predictive models of AD activity have no notion of uncertainty, leading to biased, overconfident, or misleading predictions. While neural networks do not inherently represent uncertainty, deep ensembles [35, 36] provide a scalable approach for quantifying prediction uncertainty [35]. Machine-based uncertainty can inform the optimization of predictive models in evolutionary studies by guiding the sampling of non-model organisms to build a more comprehensive training dataset.

Uncertainty can arise from inherent noise in the assay (aleatoric uncertainty) or from limited sampling within the dataset used to train the surrogate model (epistemic uncertainty). We integrated machine-based uncertainty into ADhunter to quantify epistemic uncertainty through a deep ensemble approach. Furthermore, model ensembling improves predictive performance by combining the strengths of multiple regressors, which reduces individual model biases. As expected, the model ensemble outperformed a single model on a held-out test dataset (Fig. S6). In particular, performance improves as a function of ensemble size and we selected an ensemble with 20 models (Pearson r = 0.775; RMSE = 0.632) for computational tractability. We simulated out-of-distribution inference by performing spectral clustering of the training dataset (Table S5). When evaluated on a held-out test cluster, TADA (Pearson r = 0.381; RMSE = 1.021) underperforms relative to ADhunter (Pearson r = 0.512; RMSE = 0.826). A single instance of ADhunter with neural encodings outperforms one-hot encodings, and ensembling further improves out-of-distribution performance. These results indicate that the deep ensemble improves ADhunter performance and generalizability relative to the state-of-the-art AD prediction model for evolutionary studies.

### Evolutionary sampling of transcriptional activators using machine-based uncertainty

We sought to further improve ADhunter for use in evolutionary studies with active learning from sequences in non-model genomes. Active learning involves performing inference on a design space, functional characterization of selected sequences defined by an acquisition function and updating the model on the active learning dataset [37]. This framework has successfully been used in protein engineering applications, such as in directed evolution [38], where the objective is to optimize a property of interest [39, 40]. In comparison to structured proteins, we have limited knowledge of the activity landscape for disordered proteins, such as transcriptional activators. For studying gene regulation, our objective is to minimize prediction error (i.e., the difference between empirical and predicted activity). In place of mutagenesis libraries for protein engineering [6, 40], we created an active learning library from the sequence space sampled by evolution.

Next, we performed *in silico* discovery of ADs from non-model genomes to make ADhunter robust across fungal evolutionary space. From the MycoCosm collection [41] we obtained 2,400 fungal genomes totaling 87,542,943 proteins. Sequences were deduplicated, resulting in 9,395,825 unique proteins. We clustered sequences with 90% identity to retain maximum diversity of protein space. The 7,842,516 representative sequences from each cluster were sliced into 53 AA tiles with 10 AA stride, totaling 72,057,539 tiles (Fig. 2A). ADhunter predicted the activity and associated uncertainty for each tile (Fig. 2B). There was high uncertainty at both ends of the range of predicted activity. Since only a small subset of proteins activate gene expression, this may account for the greater variance in uncertainty among sequences with low predicted activity compared to those with high predicted activity. Given that most proteins lack an AD, this likely explains the low median predicted activity across fungal protein space. In contrast to previously characterized sequences from the model fungus, the evaluated sequences encompass both unicellular and multicellular fungi with diverse morphologies and ecological niches.

**Figure 2.**
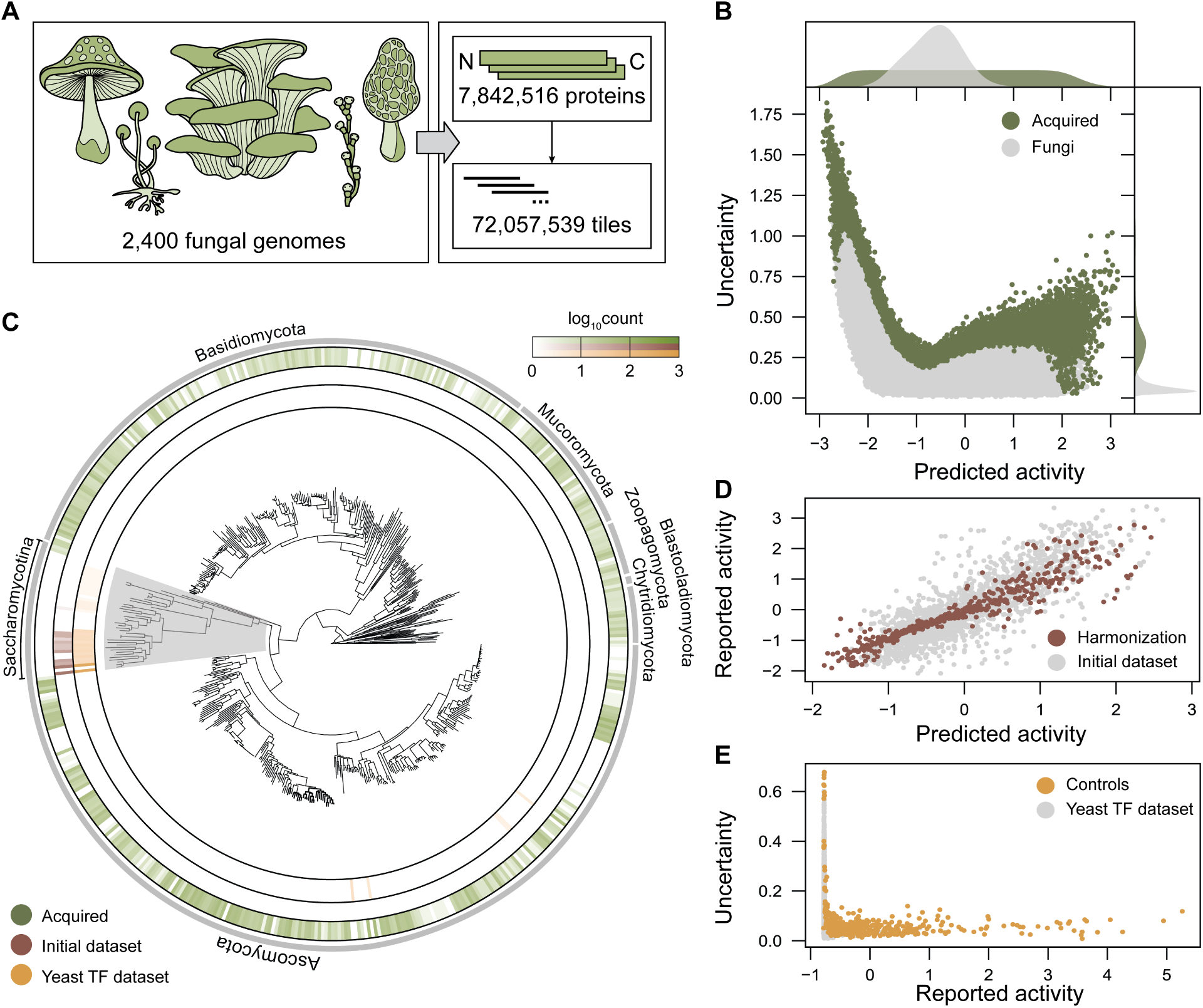
Active learning enables comprehensive exploration of fungal evolutionary space. (A) The MycoCosm collection consists of 9,395,825 unique proteins from 2,400 fungi. Clustering by 90% sequence identity resulted in 7,842,516 proteins that were tiled to perform inference with ADhunter. (B) A total of 8,935 tiles with the highest uncertainty across the range of predicted activity were selected for downstream functional characterization. The marginal histograms indicate that the median sequence exhibits negligible predicted activity and low uncertainty. ADhunter identifies a subset of sequences spanning the range of predicted activity with high uncertainty for improving model generalizability. (C) Test tiles from evolutionary space (green) represent the diversity of 1,050 non-model fungal genomes compared to 21 fungal genomes from the initial dataset (brown) and 69 fungal genomes from Sanborn et al. (yellow). Uncertainty sampling avoided sequences from the *Saccharomycotina* subdivision (grey), which contains the model fungus *S. cerevisiae*. (D) Harmonization tiles were selected from the initial dataset with the minimum error across the range of predicted activity to map the initial dataset to the active learning library. (E) Control tiles from the yeast TF dataset were selected with the highest uncertainty across the range of reported activity.

We used quantile-balanced uncertainty sampling as our acquisition function to select 8,935 sequences with maximum uncertainty across the range of predicted activity from the 2,400 fungal genomes. The acquired sequences are represented in 1,050 fungal genomes (43.8%) compared to 21 fungal genomes (0.00875%) from the initial dataset and 69 fungal genomes (0.0288%) from the yeast TF dataset [15] (Fig. 2C). Once harmonized, a process that maps the new dataset onto the same distribution of empirical activity as the initial dataset, the updated dataset will represent the diversity from 1,071 fungal genomes (44.6%). Interestingly, acquired sequences with high uncertainty originate from all clades except *Saccharomycotina*, which is sampled by the initial dataset and contain the model fungus, *S. cerevisiae*. Furthermore, the sequence diversity sampled across fungal protein space is much broader than existing datasets as shown by sequence identity clustering (Fig. S7). The sequence diversity of other datasets drops precipitously at 75% identity. Therefore, yeast and plant TFs sample a narrow sequence space relative to a function agnostic search across evolved proteins. In addition to the tiles from non-model genomes, we added control tiles and tiles that harmonize the initial and active learning datasets. For dataset harmonization, 437 tiles from the held-out test dataset were selected with the smallest difference across the range of experimental and predicted AD activity (Fig. 2D). As controls, the same acquisition function as the evolutionary sampling was used to select a subset of 464 tiles with the highest uncertainty that span the range of empirical activity from the yeast TF dataset (Fig. 2E). Our active learning library provides comprehensive exploration of fungal evolutionary space for functional characterization of diverse ADs from non-model fungi, including deeper sampling within *Ascomycota* and the first high-throughput characterization across fungal divisions.

Completing the active learning cycle requires functional characterization of the new library, harmonizing the initial and new datasets, then retraining ADhunter on the harmonized dataset. We used a previously developed high-throughput assay [11] to quantify AD activity (Fig. 3A). For each tested tile there was a median of 12 DNA barcodes (Fig. 3B). We recovered 379 of 464 control tiles (81.7%), 404 of 437 harmonization tiles (92.4%) and 7,681 of 8,935 non-model fungal tiles (86.0%) (Fig. 3C). As expected, the control tiles spanned the range of empirical activity with high correlation to the activity reported in Sanborn et al. (Pearson r = 0.695; RMSE = 0.841; Fig. 3D). Harmonization tiles in the active learning dataset and the initial dataset had high correlation (Pearson r = 0.934; RMSE = 0.274; Fig. 3E). Overall, the test tiles from non-model fungi had lower activity relative to the initial dataset. Interestingly, there was poor correlation between the empirical and predicted activity for test tiles from non-model fungi (Pearson r = 0.178; RMSE = 1.15). These results indicate that there are novel sequence-to-function relationships within proteins from non-model fungi in the active learning dataset that were not in the initial dataset.

**Figure 3.**
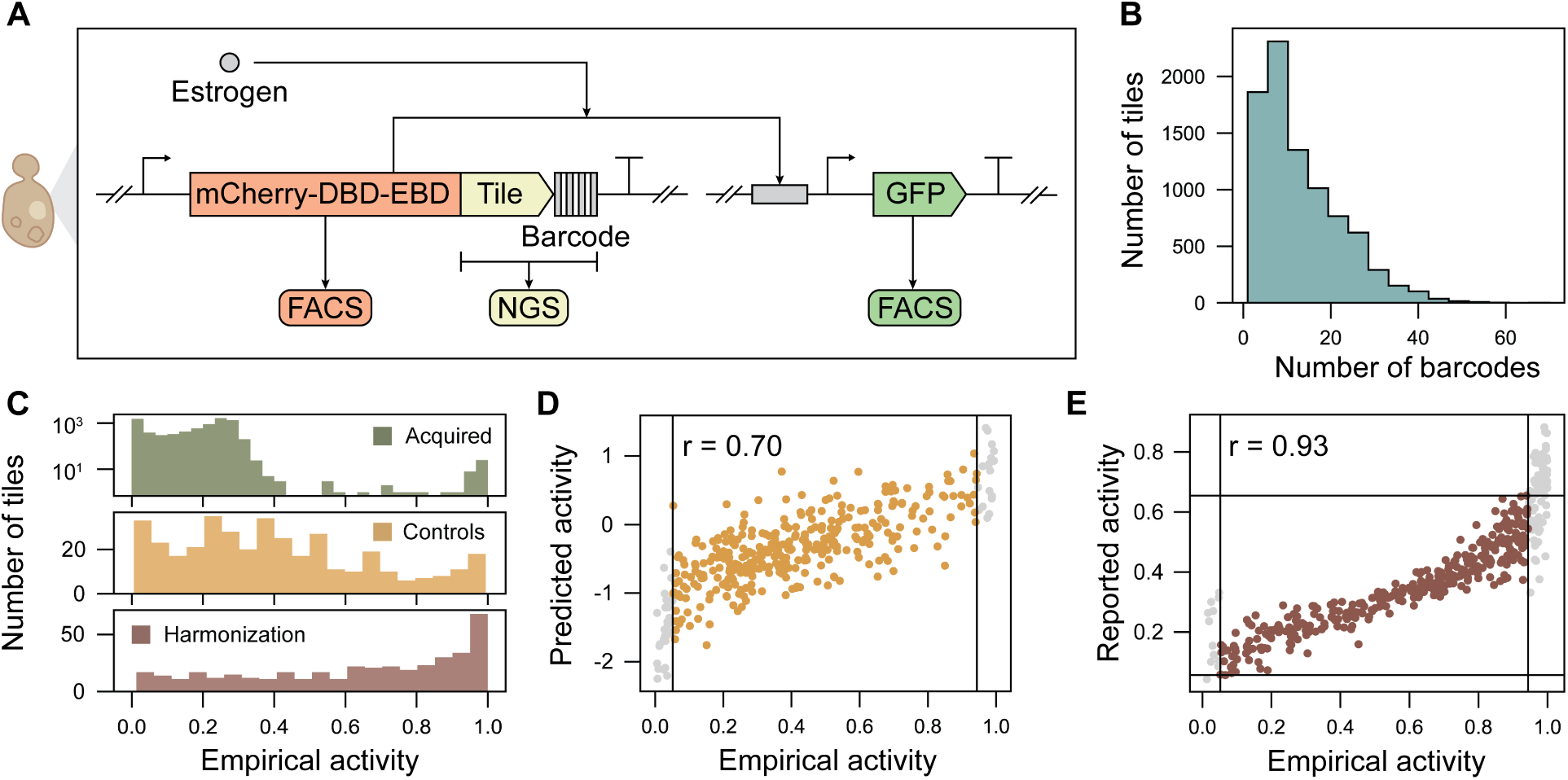
Functional characterization of transcriptional activators from non-model fungi. (A) Test sequences are cloned into a transcriptional unit containing a constitutive promoter driving expression of a red fluorescent protein (mCherry) fused to a DNA-binding domain (DBD), an estrogen-binding domain (EBD), and the test tile, followed by a 14 nt DNA barcode and terminator. Estrogen treatment induces translocation of the synthetic TF from the cytoplasm to the nucleus, thereby activating expression of a green fluorescent protein (GFP) reporter. Empirical activity for each test sequence is measured by fluorescence-activated cell sorting (FACS) of the GFP:mCherry ratio and next-generation sequencing (NGS) of the DNA barcode from sorted cells. (B) The active learning library had a median of 12 DNA barcodes per tile, and AD activity was averaged across the barcodes for each corresponding AD tile. (C) The active learning library covered the full range of the GFP:mCherry signal, with the high uncertainty tiles from non-model fungi (green) primarily appearing in the lower range of empirical activity. The control tiles (yellow) and harmonization tiles (brown) spanned the range of empirical activity. (D) Control tiles had high correlation to the predicted activity where tiles in grey are outside the linear range of the assay. (E) Harmonization tiles selected had high correlation across studies, enabling the initial and active learning datasets to be mapped onto the same activity distribution.

The active learning dataset had exact matches to 17,992 proteins across fungal divisions (Fig. 4A). Sequences capable of activating gene expression have previously been identified in proteins that are not typi-cally involved in transcriptional regulation [16]. Characterized tiles map to 1,020 proteins involved in cellular processes and 2,551 involved in metabolism. Across fungal divisions, 3,416 are poorly characterized. The acquired tiles have exact matches to 2,210 proteins involved in information processing, where 1,101 proteins have annotated DNA-binding domains (DBDs) (Fig. 4B). The majority of proteins with DBDs correspond to zinc cluster TFs, which are involved in transcriptional control of secondary metabolism and pathogenesis and often contain a C-terminal AD. Functionally characterized tiles in the active learning dataset map to 7,250 proteins in *Ascomycota*, which represents the largest division of sequenced fungal genomes. These sac fungi range from unicellular yeast to large, multicellular molds and morels that produce a range of natural products. Biosynthetic gene clusters in these organisms are often silent under laboratory conditions. ADhunter can facilitate their programmable expression by identifying cryptic transcriptional activators. In addition to *Ascomycota*, there are tiles that map to 1,355 proteins in *Basidiomycota*, which are being engineered for food and sustainability applications. For example, edible mushrooms require activation of heterologous genes for novel flavor profiles. Rusts that spread crop diseases could be inactivated by improved mapping of transcriptional activators controlling pathogenesis genes. Evolutionary sampling provides the first systematic annotation for mapping genetic regulation in understudied branches of life.

**Figure 4.**
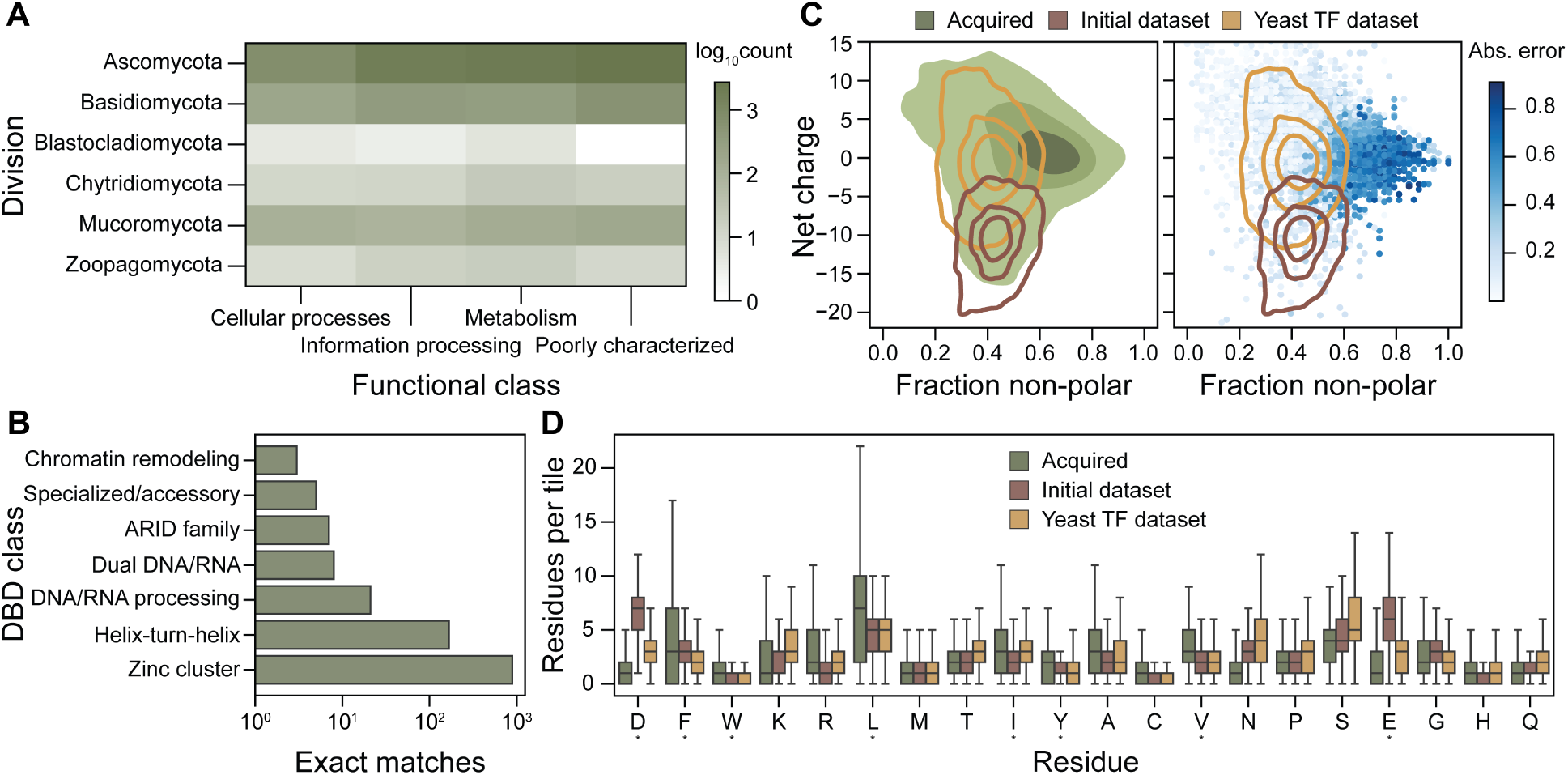
Non-model fungal proteins exhibit out-of-distribution sequence features. (A) The active learning library contains tiles with exact matches to proteins that span taxonomical divisions and functional classes, especially 3,416 that are poorly characterized or have unknown function. (B) Tiles from the active learning library have exact matches 1,101 TFs, the majority of which belong to the Zinc cluster DNAbinding domain (DBD) class, which are regulators of metabolism, stress responses, and natural product biosynthesis. (C) Tiles acquired from non-model fungal genomes exhibit more diverse sequence properties, particularly in net charge and fraction of non-polar residues-compared to datasets from model fungi. The initial model fails to accurately predict the activity of tiles with sequence features that lie outside the range observed in yeast datasets. (D) The number of residues per tile of the active learning library (green) to the initial dataset (brown) and the yeast TF dataset (yellow). Residues from the acidic exposure model are noted with an asterisk.

The distribution of net charge versus fraction of non-polar residues in the initial dataset and yeast TF dataset show that prediction error originates outside the sampled distribution (Fig. 4C). The active learning dataset acquired sequences that span both distributions with a higher overall fraction of non-polar residues. In particular, the acquired tiles are non-acidic as well as leucine- and phenylalanine-enriched relative to datasets from the model yeast (Fig. 4D). The strongest tiles from non-model fungi, as well as those with the largest prediction error, contain a higher fraction of leucine and phenylalanine residues compared to previously characterized datasets. Tiles above the median absolute prediction error are found in 7,236 proteins across 849 fungal genomes. Therefore, adding these sequences with high uncertainty to the training dataset should enable ADhunter to identify ADs from a larger sequence space relative to the initial dataset.

### Active learning enables quantification of protein codes from non-model fungal genomes

After harmonizing the datasets, we evaluated the improvement in performance from active learning on a new held-out test dataset with sequences from both the initial and newly acquired dataset. The held-out test dataset represents the diversity of 406 fungal genomes (Fig. 5A). ADhunter trained on the initial dataset was able to perform well on the test sequences from yeast, but there was a clear subset of sequences with high prediction error and uncertainty from non-model fungi (Pearson r = 0.541; RMSE = 0.963). In particular, prediction error and uncertainty was highest outside sequences from *Saccharomycotina*. However, ADhunter trained on the harmonized dataset was able to identify these patterns and achieved much lower prediction error and uncertainty across the held-out test dataset (Pearson r = 0.824; RMSE = 0.570). These results demonstrate that active learning reduced uncertainty in ADhunter and identified novel sequence-to-function relationships of ADs from a diverse sampling of evolutionary space. Therefore, active learning improved ADhunter’s ability to generalize across fungal evolutionary space, enabling deeper insights for evolutionary studies of gene regulation.

**Figure 5.**
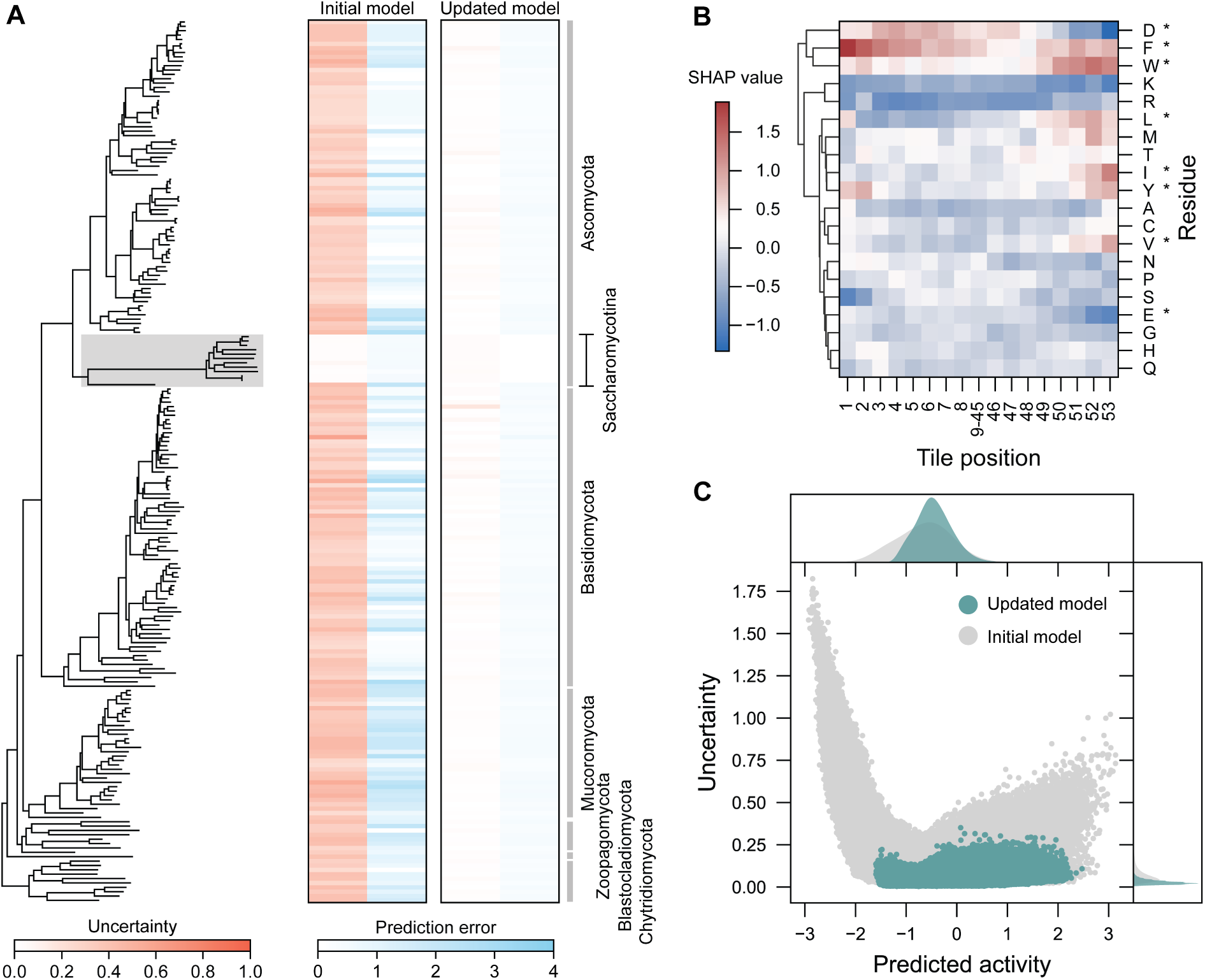
Active learning enables accurate quantification of sequence-to-function relationships across evolutionary space. (A) ADhunter trained on the initial dataset was unable to detect sequenceto-function relationships from non-model organisms. Updating ADhunter with a subset of the active learning dataset and evaluating on a held-out test dataset shows reduced prediction error and uncertainty. ADhunter trained on the initial dataset had high uncertainty outside the Saccharomycotina subdivision (grey) compared to the lower uncertainty across fungal protein space of the updated model. (B) Model interpretability analysis identifies sequence contributions governing AD activity that align with experimental observations and biophysical models. Residues from the acidic exposure model are noted with an asterisk. The majority of functional determinants are within the first and last 8 residues. (C) Performing inference using the updated model across fungal protein space reduces uncertainty, enabling ADhunter to be used in evolutionary studies.

We further evaluated ADhunter with respect to the state-of-the-art by partitioning performance contributions attributed to the dataset composition versus the prediction task (Table S6). Comparison across models is complicated by differences in AD sequence lengths. To assess the role of AD sequence length, we retrained ADhunter and TADA separately on the 40 AA from Morffy et al. or the 53 AA harmonized dataset. ADhunter achieved superior performance relative to TADA when evaluated on both held-out test datasets. Comparison across models is further complicated by differences in model optimization objectives. ADhunter minimizes a regression objective (i.e., mean squared error loss) whereas TADA minimizes a classification objective (i.e., focal loss). To assess classification performance, we binarized the predicted activity by ADhunter as described in Morffy et al. and found that ADhunter outperforms TADA. To assess regression performance, we used the TADA score as the predicted activity and found that ADhunter provides more quantitative predictions. Overall, these findings further demonstrate that ADhunter achieves state-of-the-art performance across datasets and prediction tasks.

Next, we examined how ADhunter quantifies activity from protein sequences relative to an established biophysical model of ADs. Using SHapley Additive exPlanations (SHAP) [42], we analyzed sequence feature contributions and found that functional determinants of AD activity tend to cluster at the Nand C-termini (Fig. 5B). Sequence features in the middle of the tile exhibit more consistent, residue-level contributions (Fig. S8). Consistent with the acidic exposure model [19], Asp, Phe, and Trp most strongly increased predicted activity, whereas Lys and Arg contribute most to its reduction. We also observed that all residues important in the acidic exposure model have contributions that increased predicted activity at the C-terminus, except for Asp and Glu which decrease predicted activity. Interestingly, Val and Glu have a much smaller impact on predicted activity compared to other previously described residues. As experimentally observed [16, 43], Leu, Met, and Trp have a significant impact on AD activity at the C-terminus. Model interpretability analysis of ADhunter is consistent with the acidic exposure model, suggesting that ADhunter’s predictions align with known AD sequence characteristics. These results complement the model interpretability performed by Morffy et al. by providing single-residue resolution of sequence contributions to AD activity.

We revisited fungal protein space using ADhunter trained on the harmonized dataset and found an overall reduction in uncertainty and a smaller range of predicted activity with respect to ADhunter trained on the initial dataset (Fig. 5C). The reduced uncertainty in characterized tiles from non-model fungi and across fungal protein space suggest that active learning improved model generalizability across evolutionary space. ADhunter enables accurate quantification of ADs across fungal evolutionary space and our framework can be extended to study biological properties across underexplored branches of life.

## Discussion

Biological discovery and design are increasingly being guided by surrogate models trained on data from high-throughput technologies. While large-scale experimentation is often performed in model species, acquisition of functional measurements (i.e., ground truth labels) should not be limited to sequences from these organisms. In this study, we demonstrate how active learning enables comprehensive exploration of fungal evolutionary space, thereby enhancing model generalizability for predicting biological properties. We use machine-guided exploration to traverse the sequence landscape of transcriptional activators in fungi, a largely uncharacterized branch of life that represent 26.9% of all eukaryotic reference genomes. Predicting properties of intrinsically disordered proteins remains a significant challenge, even with the most advanced computational models. We present a framework that fine-tunes pretrained neural encodings to accurately predict transcriptional activation of disordered proteins across evolutionary space. By integrating high-throughput technologies with advances in machine learning, this framework extends beyond transcriptional activators to enable robust functional genomics models that quantify biological properties at an evolutionary scale.

ADhunter outperforms state-of-the-art models in classification and quantitation of ADs for use in evolutionary studies. We optimize model performance by replacing binary protein representations with continuous neural encodings and deep ensembling. These features also improve model generalizability, as demonstrated by spectral clustering analysis. While design spaces in protein engineering often center on mutagenesis libraries, our approach leverages the rich diversity of sequencing data to explore sequences across non-model species. Evolved proteins are highly diverse and enriched for functional sequences, offering a more expansive foundation for robust quantification of biological properties using predictive models.

A key feature of ADhunter is its integration of machine-based uncertainty. By prioritizing sequences with high uncertainty across the range of predicted activity, we focus experimental efforts on maximally informative samples. We performed functional characterization for a library of diverse sequences, which significantly improved ADhunter’s ability to generalize outside *Saccharmoycotina* and across fungal divisions. This approach enables quantification of underrepresented sequence-to-function relationships from non-model organisms compared to existing models that tend to identify overrepresented patterns. Our results show how ADhunter enables dissection of the evolutionary trajectory and neofunctionalization of transcriptional activators in natural genetic circuits. In the context of fungal biology, ADhunter can be used to study functional evolution in gene regulatory networks underlying niche expansion and transcriptional control of natural product pathways. For fungal engineering, ADhunter can guide the design of TFs for inducible promoters with fine-tuned expression properties. Transcriptional activation of synthetic programs is especially important as interest grows in engineering non-model fungi for sustainability and food applications.

As predictive models trained on high-throughput datasets continue to guide biological discovery and design, addressing prediction biases from the training dataset composition is crucial. Our work highlights the importance of expanding functional characterization beyond model organisms to include sequences from non-model species. By leveraging advances in high-throughput molecular technologies and machine learning, we can accelerate the study of underexplored branches of life to identify universal principles of living systems and reprogram organisms for novel purposes.

## Methods

### Code and sequencing availability

All code related to this study are publicly available on GitHub (github.com/shih-lab/ADhunter). The base model of ADhunter using binary encodings is publicly available on GitHub (github.com/staller-lab/adhunter). Raw sequencing reads are publicly available through the NCBI SRA Database under BioProject accession PRJNA1183837. DNA oligonucleotides are listed in the Supplementary Data.

### Plasmid library assembly

Protein sequences from the MycoCosm collection were synthesized by Twist Biosciences as tiles in a DNA oligonucleotide pool. Each tile is flanked by a conserved upstream (GCGGGCTCTACTTCATCGGCTAGC) and downstream (TGATAACTAGCTGAGGGCCCG) sequence for Gibson cloning into pMVS142 (Addgene #99049). The DNA oligonucleotide pool was used in a PCR using Q5 polymerase (New England Biolabs, USA) with primers LC3.P1_Lib_Hom_up and YL_randBCs_R3 to add homology arms and a random 14 nt barcode. The PCR was cleaned using DNA Clean and Concentrator 5 (Zymo Research, USA). pMVS142 was linearized using NheI, AscI, and PacI (New England Biolabs, USA). The barcoded tiles and linearized plasmid were assembled using HiFi DNA Assembly (New England Biolabs, USA). Assemblies were electroporated into 10-beta Competent *E. coli* (New England Biolabs, USA) and recovered for 20 minutes at 37 °C. Approximately 2M colonies were recovered then isolated using the Plasmid Maxi Kit (Qiagen, USA).

### Yeast library construction

The plasmid library was transformed into the URA3 locus of strain DHY211 using a standard lithium acetate method. The plasmid library was digested with SalI, EcoRI, and PacI (New England Biolabs, USA) to reduce background plasmids missing a barcoded tile. Homology arms were amplified from pMVS295 and pMVS296 for integration into the URA3 locus using primers YP18/CP19.P6 and YP7/YP19, respectively. The yeast was treated for 30 minutes at 30 °C then 60 minutes at 42 °C. The transformed library was plated on YPD, followed by overnight incubation at 30 °C, and replica plating onto SC+G418+5-FOA. The transformants were mated with a FY5 strain containing the reporter construct inserted into the YBR032 ORF. Diploids were selected on YPD+G418(200 µg/mL)+NAT(100 µg/mL). The yeast transformants were mated in batches then pooled and stored.

### Fluorescence-activated cell sorting and sequencing

The yeast library was grown overnight in SC+G418+NAT at 30 °C prior to sorting. Cultures were diluted (1:5) into SC+beta-estradiol (1uM) and incubated for 4 hours at 30 °C. Yeast libraries were sorted using an Aria Fusion Flow Cytometer (BD Biosciences, USA) at the UC Berkeley Flow Cytometry core facility. As a negative control, the parent strain with the reporter and no barcoded tile was used to calibrate baseline values in the GFP:mCherry channel. A positive control strain expressing the strong VP16 AD was used to calibrate the high end of the GFP:mCherry ratio channel. A total of 500,000 cells were sorted into each of 8 bins across GFP:mCherry ratios, with each bin representing 10% of the population. Sorted cells were grown overnight in SC at 30°C. Genomic DNA was extracted using the Wizard Genomic DNA Purification Kit (Promega, USA). DNA barcodes were amplified by PCR using primers CP21.P14 and CP17.P12 for 20 cycles. Phasing nucleotides were added by PCR using primers SL5.F18 through SL5.F21 and SL5.R1 through SL5.R4 for 10 cycles. Sequencing indices were added by PCR using primers MVS_0001_I1 through MVS_0024_I1 and MVS_0001_I2 through MVS_0024_I2 for 6 cycles. Sequencing libraries were cleaned using DNA Clean and Concentrator 5 (Zymo Research, USA) to remove residual primers. The UC Berkeley Sequencing Facility normalized and sequenced libraries on an Illumina NovaSeq X 10B with 2x150 bp paired-end reads.

### Model implementation

ADhunter consists of a convolutional layer, a series of residual blocks, a pooling layer, and a fully-connected output layer. Each residual block contains two convolutional layers with batch normalization and ReLU activation. ADhunter optimizes mean squared error loss using Adam and outputs evaluation metrics for root mean squared error, Pearson correlation, and Spearman correlation. To prevent overfitting, we added early stopping if the validation loss did not improve after 5 epochs. Labeled datasets were preprocessed by removing duplicate entries and z-score normalizing the activity values where 80% was used for training, 10% for validation, and 10% for testing. Since AD activity is continuous, the value was binarized with respect to the median activity to enable stratified shuffling of the data. Protein sequences and activation measurements from Hummel et al. were used for the initial training dataset. ESM embeddings were obtained as described on the ESM GitHub repository (github.com/facebookresearch/esm).

### Model interpretability analysis

The DeepExplainer module from SHAP was used to evaluate model interpretability. For each model in the ensemble, background samples from 1,000 randomly selected training data were used and evaluated on the held-out test dataset. SHAP values were averaged over the encodings over the full sequences.

### MycoCosm analysis

Protein sequences were obtained from MycoCosm via the Joint Genome Institute. Protein sequences were deduplicated then clustered by sequence identity using CD-HIT[44]. The test tiles from the MycoCosm collection were codon optimized for expression in *S. cerevisiae* whereas the original codons were used from dataset harmonization and control tiles.

### Sequencing analysis

Raw sequencing reads were demultiplexed using bcl2fastq v2.19.0, trimmed using trimmomatic[45] v0.39, and assembled using PANDAseq[46] v2.11. Only reads with a perfect match to a tile in the library were retained. Tile and barcode sequences were extracted using a custom regular expression matching conserved regions. For each set of eight sorted samples, reads were normalized by the total number of reads in each bin. For each tile, counts were normalized across the eight sets to calculate a relative abundance. Activity scores were calculated by taking the inner product between relative abundances and with the median fluorescent value of each bin, resulting in a weighted average. Tiles with less than 50 reads were discarded. We used labtools v0.0.3 to quantify activity for each AD tile (github.com/staller-lab/labtools). The activity score for a given tile was aggregated by taking the mean activity across all barcodes.

### Dataset harmonization

The FACS saturates at high and low GFP:mCherry signal. We determined the linear range of the assay by maximizing the Pearson correlation between harmonization tiles in the initial and new dataset then validating the thresholds on the Pearson correlation of the control tiles. To harmonize the two datasets we used a linear fit with Gaussian noise to map the activity values onto the same distribution. When evaluating the performance improvement of active learning, ADhunter was retrained on the harmonized dataset and a random held-out test dataset was used for evaluation. ADhunter was retrained on the entire harmonized dataset when performing inference on MycoCosm.

## Acknowledgements

Many thanks to Igor Shabalov for providing the phylogenetic tree of the fungal genomes in MycoCosm. Thank you to Ryan Friedman, Claire LeBlanc, Melvin Soriano, Pippa Richter, and members of the Shih lab for helpful comments on the work. Lucas Waldburger is funded through the National Science Foundation Graduate Research Fellowship. Max Staller is a Chan Zuckerberg Biohub – San Francisco Investigator funded by NIH grant number R35GM150813. Angelica Lam and the development of ADhunter was supported by NSF grant number 2112057. This work was part of the DOE Joint BioEnergy Institute supported by the U.S. Department of Energy, Office of Science, Office of Biological and Environmental Research, supported by the U.S. Department of Energy, Energy Efficiency and Renewable Energy, Bioenergy Technologies Office, through contract DE-AC02-05CH11231 between Lawrence Berkeley National Laboratory and the U.S. Department of Energy. The views and opinions of the authors expressed herein do not necessarily state or reflect those of the United States Government or any agency thereof. Neither the United States Government nor any agency thereof, nor any of their employees, makes any warranty, expressed or implied, or assumes any legal liability or responsibility for the accuracy, completeness, or usefulness of any information, apparatus, product, or process disclosed, or represents that its use would not infringe privately owned rights. The United States Government retains and the publisher, by accepting the article for publication, acknowledges that the United States Government retains a nonexclusive, paid-up, irrevocable, worldwide license to publish or reproduce the published form of this manuscript, or allow others to do so, for United States Government purposes. The Department of Energy will provide public access to these results of federally sponsored research in accordance with the DOE Public Access Plan.

## Contributions

Conceptualization, L.W., H.N., P.M.S., M.V.S.; Resources and supervision, P.M.S.; Methodology, L.W., H.N., A.L., M.Z., L.D.K.; Investigation, L.W., H.N., M.Z., L.K., N.L.; Writing Original Draft, L.W., P.M.S., M.V.S.; Writing Review and Editing, All authors.

## Competing Interests

J.D.K. has financial interests in Amyris, Ansa Biotechnologies, Apertor Pharma, Berkeley Yeast, BioMia, Demetrix, Lygos, Napigen, ResVita Bio, and Zero Acre Farms. P.M.S. has financial interests in BasidioBio.

## Supplementary Figures

**Figure S1.**
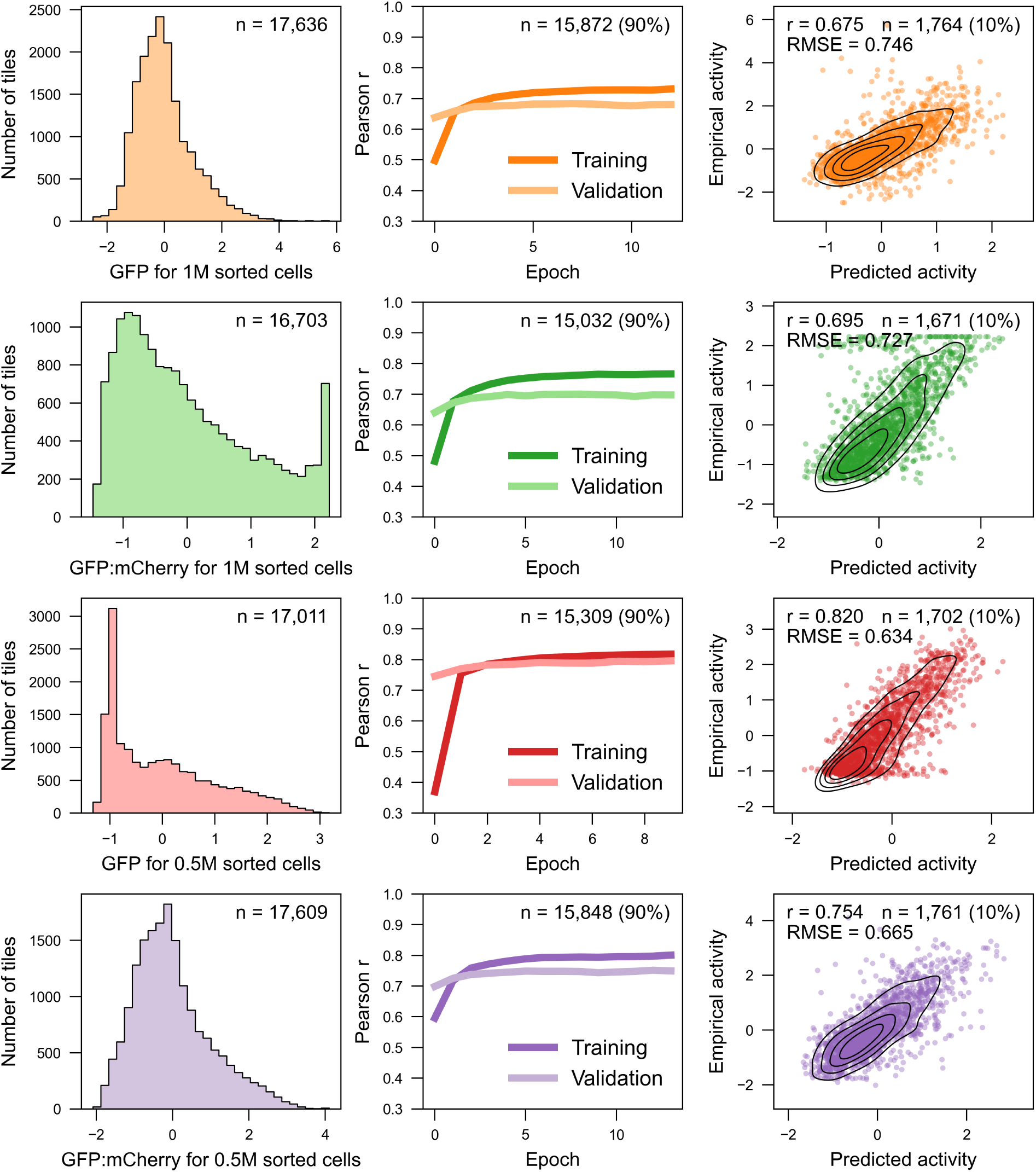
Selection of AD activity metric. The AD activity measurement was selected based on the distribution of empirical activity. We used the GFP:mCherry ratio for 500,000 sorted cells for sequence-tofunction modeling of transcriptional activators.

**Figure S2.**
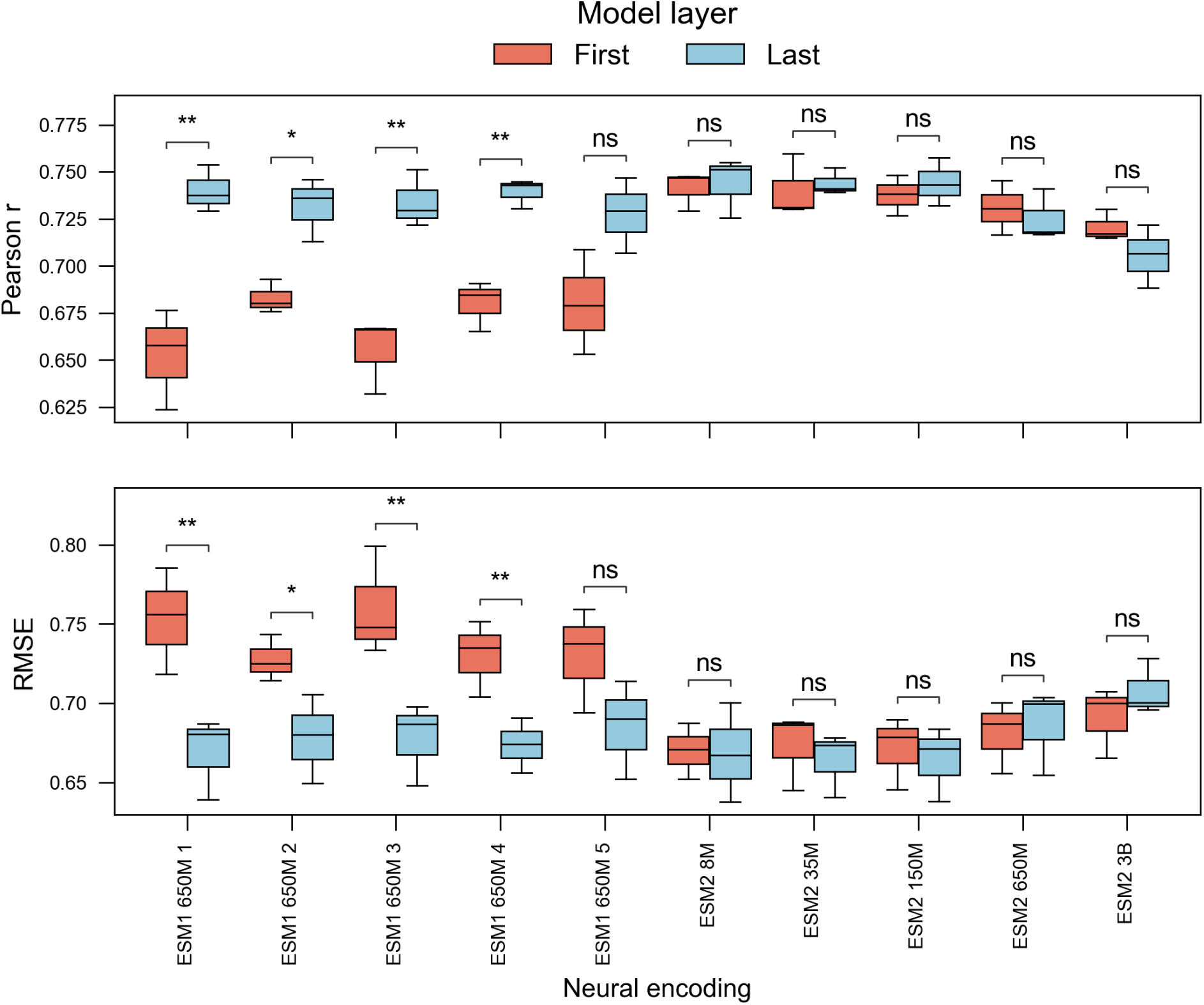
Selection of protein sequence encoding. Neural encodings slightly outperform simple encodings and were selected for predictive modeling. A two-sided Student’s t-test with Bonferroni multiple comparison correction was used to assess statistical significance between the first and last layer of each ESM model. Results show that encodings from the first layer versus the last layer of version 1 models have statistically significant differences in performance on a held-out test dataset (ns: 5e-2 < p ≤ 1; *: 1e-2 < p ≤ 5e-2; **: 1e-3 < p ≤ 1e-2; ***: 1e-4 < p ≤ 1e-3, Student’s t-test with Bonferroni multiple comparison correction). Each model and layer combination was evaluated across 3 random seeds.

**Figure S3.**
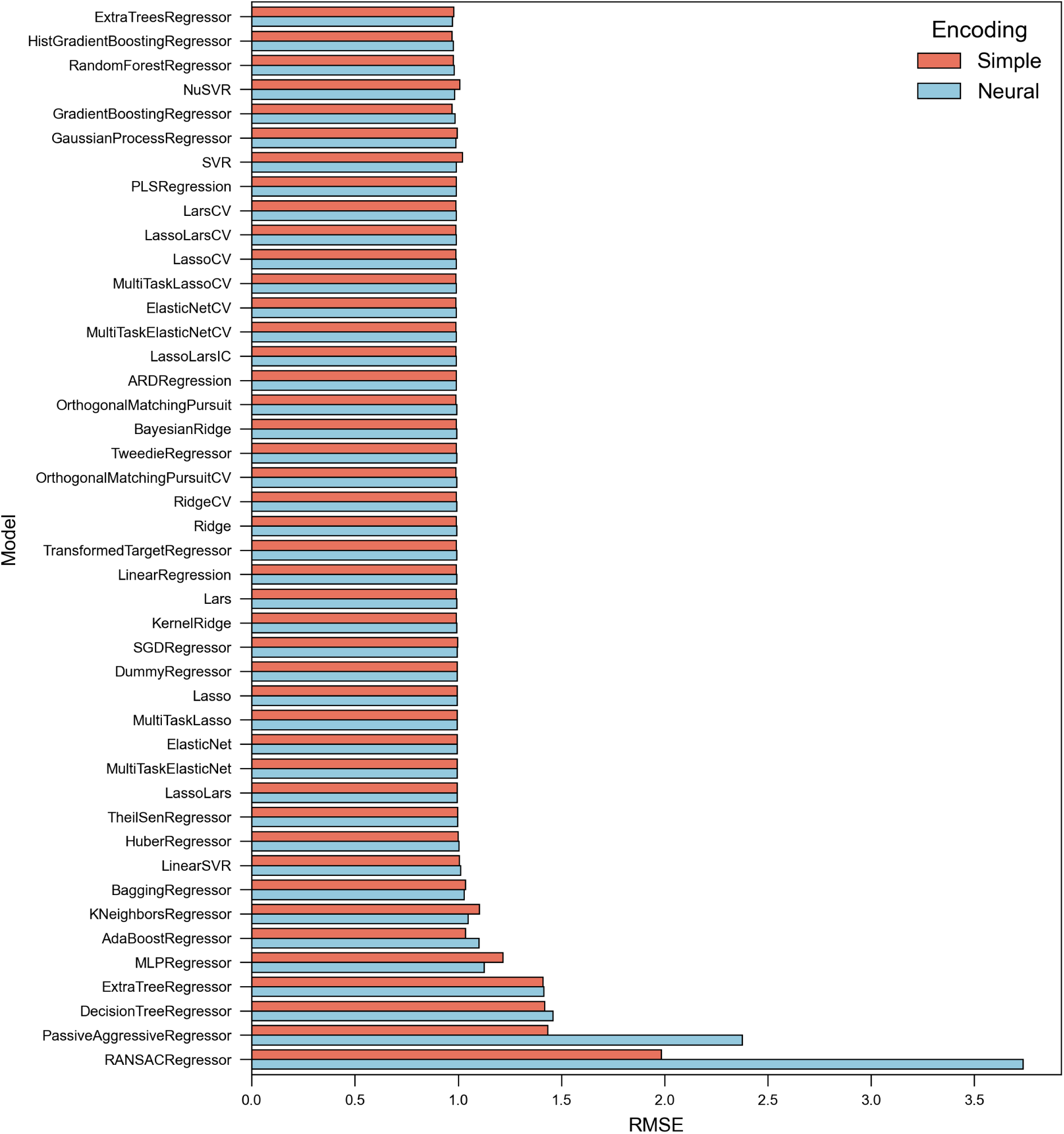
Evaluation of lightweight regressors. We evaluated 44 lightweight regressors trained with simple integer encodings or pretrained neural encodings. All models performed poorly on the held-out test dataset, leading us to consider more complex architectures for quantifying transcriptional activators.

**Figure S4.**
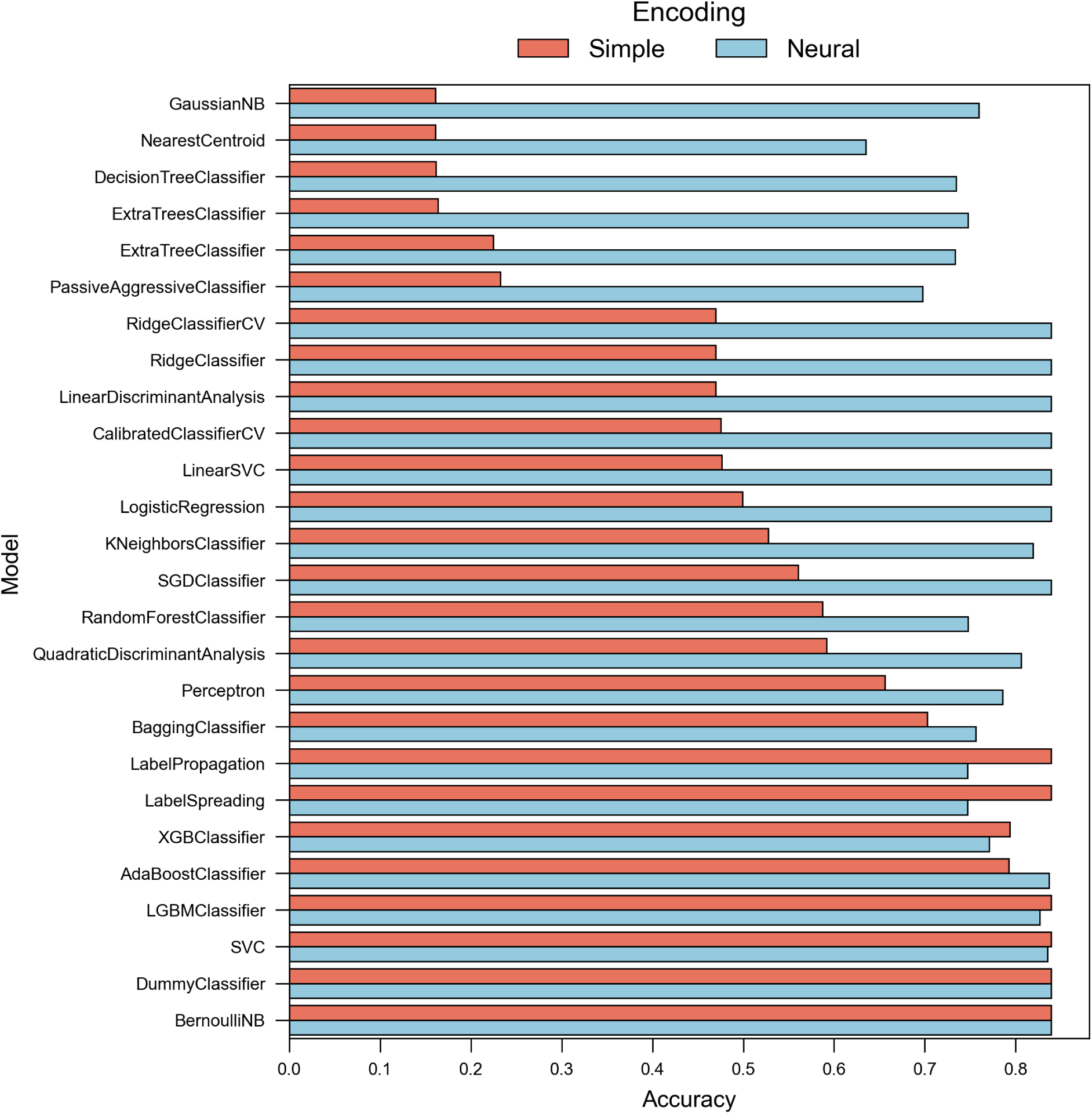
Evaluation of lightweight classifiers. Despite poor performance of the lightweight regressors, several lightweight classifiers had high accuracy performed on the binary classification task. These results indicate that the classification task has lower complexity than the regression task and that several classifiers perform better when trained with neural encodings over simple encodings.

**Figure S5.**
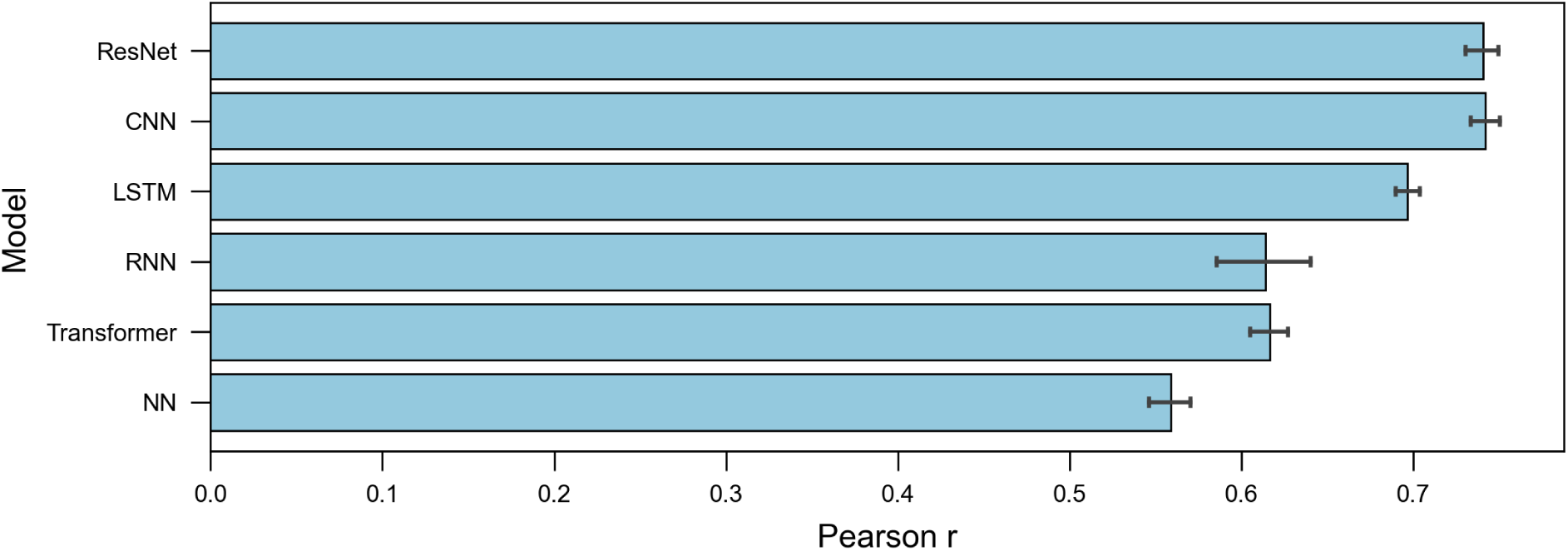
Selection of surrogate model architecture. Complex architectures outperform lightweight regressors on a held-out test dataset. Models incorporating convolutional layers, specifically a CNN and a CNN with residual connections (ResNet), achieve the highest prediction performance. Each model was evaluated across 10 random seeds.

**Figure S6.**
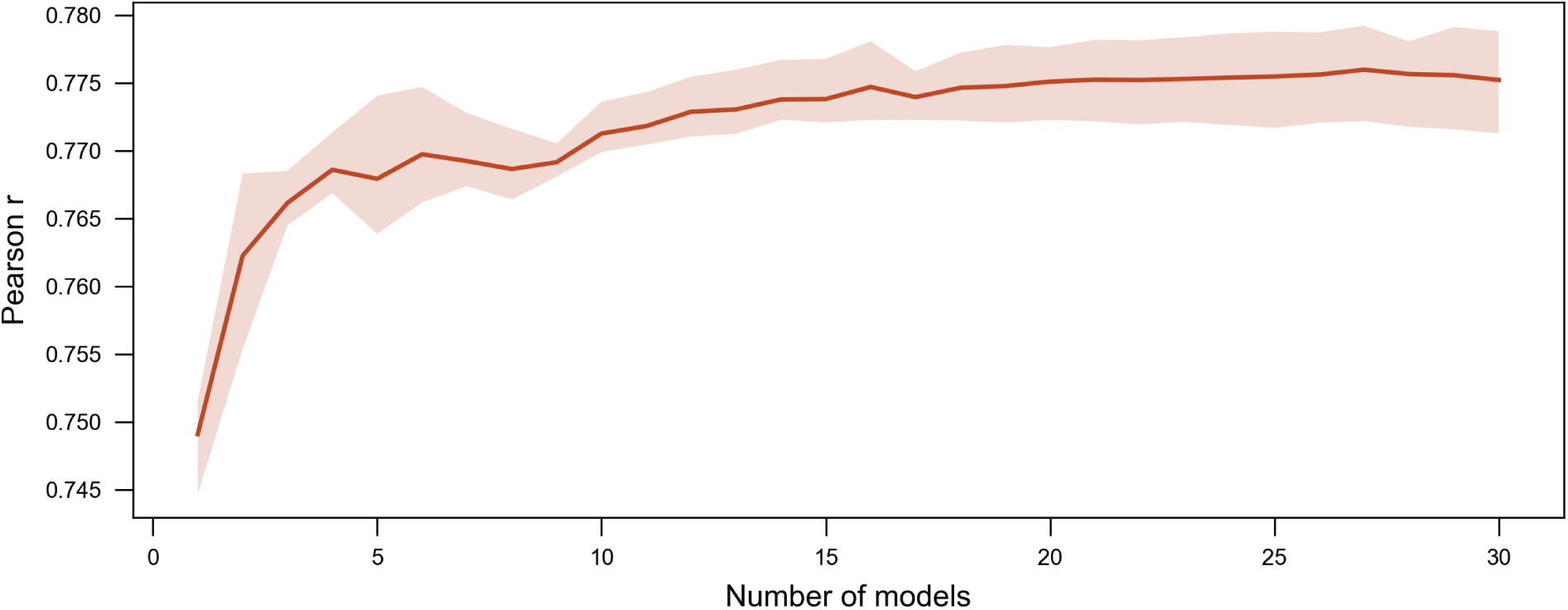
Deep ensembling of ADhunter. Improvement in ensemble model performance with respect to ensemble size on a held-out test dataset across 10 random seeds.

**Figure S7.**
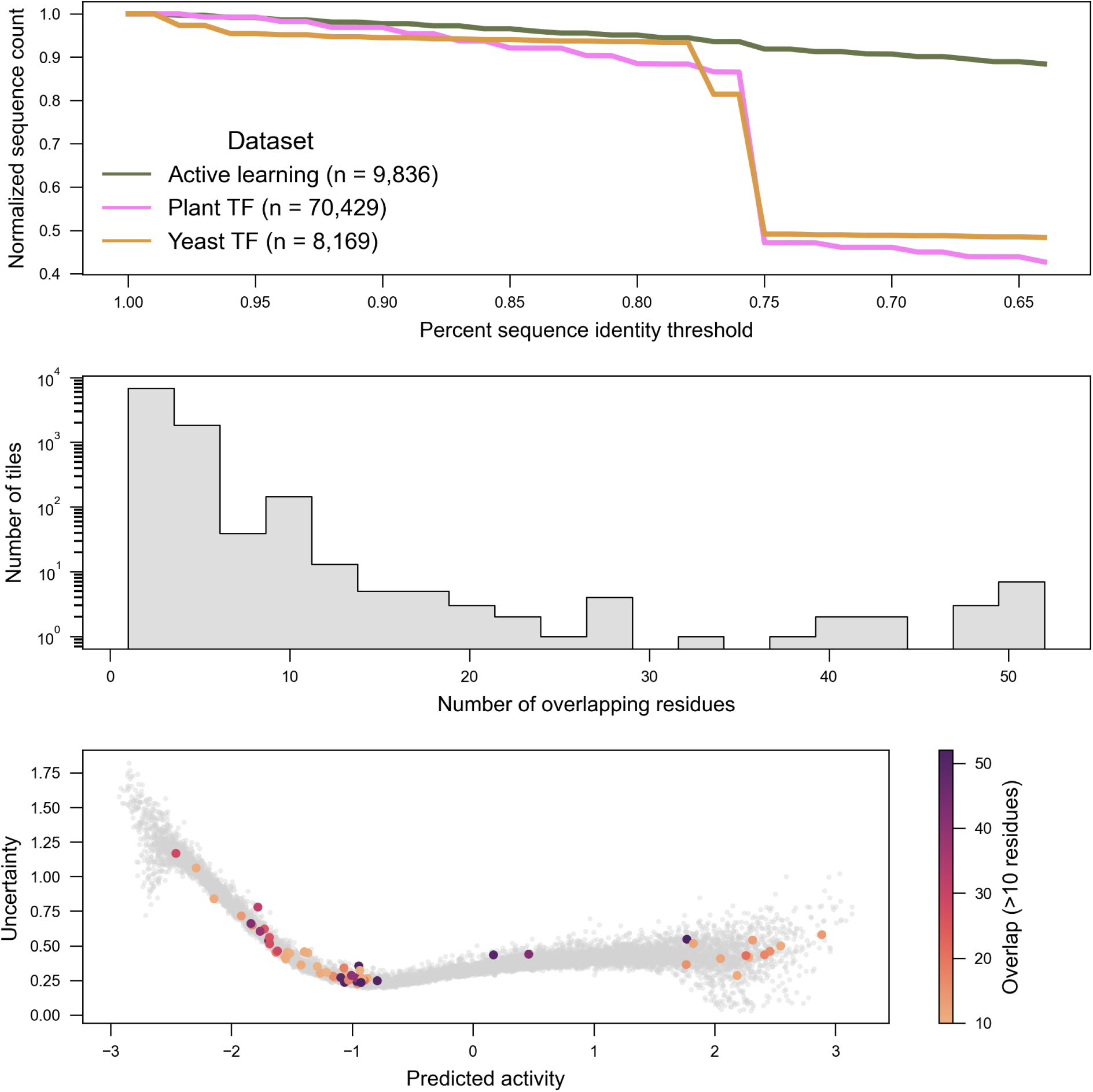
Dataset sequence identity clustering. Each dataset was clustered by sequence identity and the active learning dataset selected from the MycoCosm collection is shown to contain the most diversity. Within the test dataset, the median sequence overlap between tiles is 3 residues, where tiles with more than 10 overlapping residues are selected from across the range of predicted activity.

**Figure S8.**
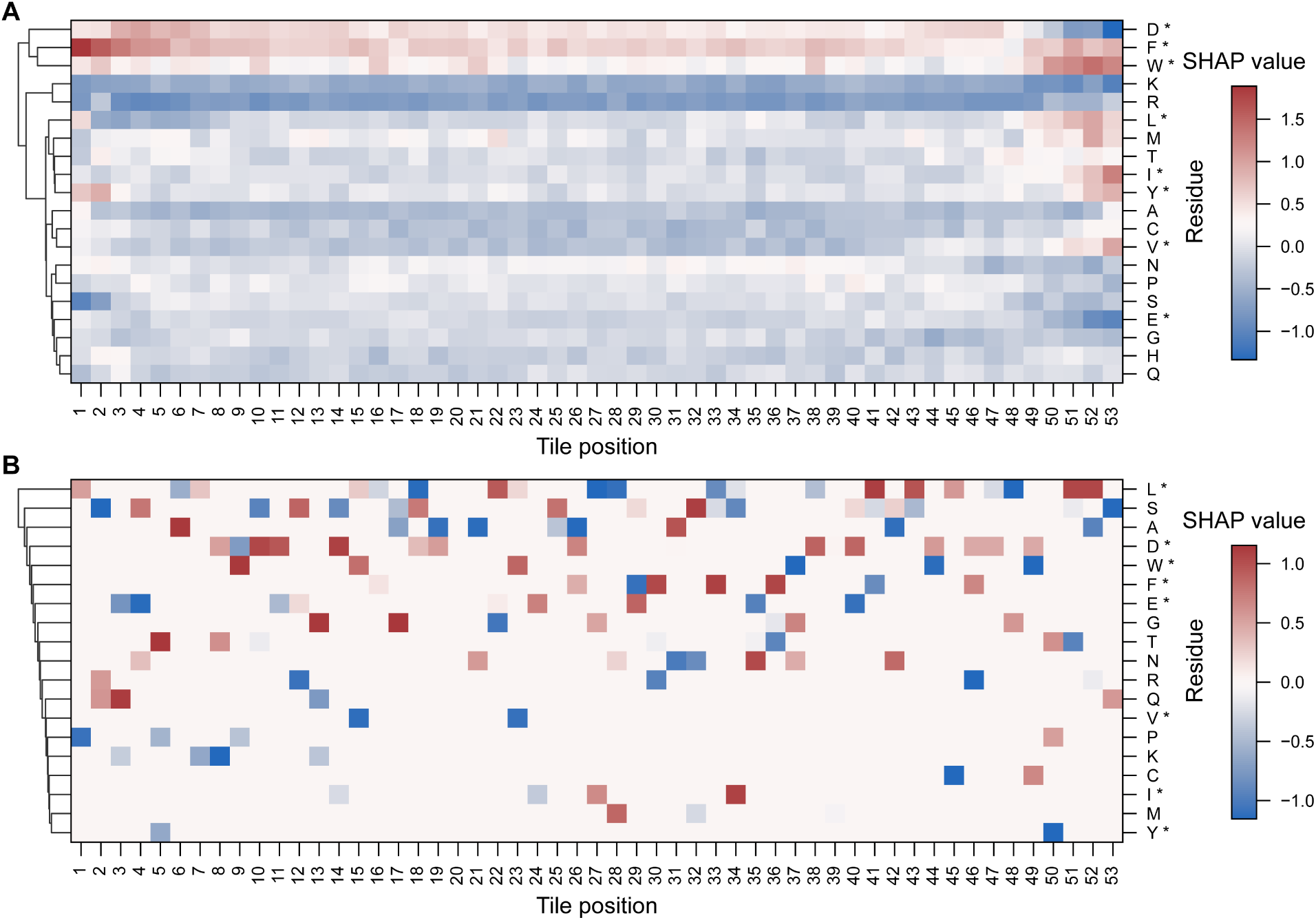
Model interpretability analysis of transcriptional activators. (A) SHAP analysis shows the majority of functional determinants are within the first and last 8 residues. (B) Strong activators (i.e., mean + 1 s.d. as defined in Morffy et al.) have very specific sequence grammar.

## Supplementary Tables

**Table S1.** Evaluation of PADDLE performance. PADDLE was evaluated on the initial dataset and achieved poor prediction performance. The model combines one-hot encodings and secondary structure predictions. However, the secondary structure predictions result in worse performance likely due to increased complexity of input features.

| Secondary Structure Included | Pearson r | RMSE |
| --- | --- | --- |
| True | 0.261 | 0.336 |
| False | 0.338 | 0.329 |

**Table S2.**
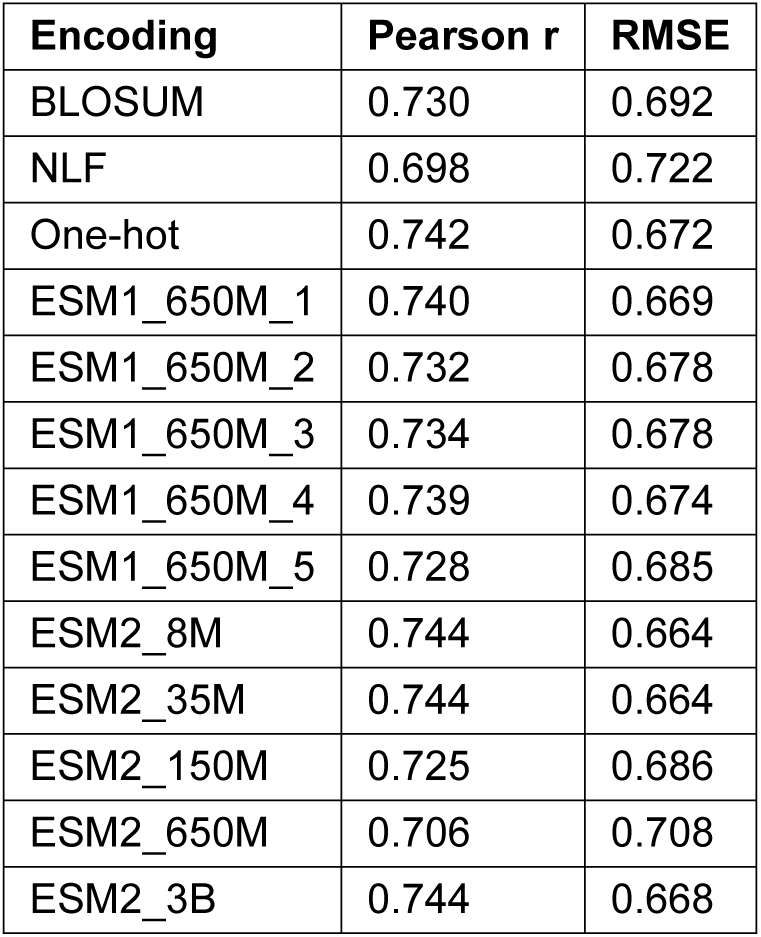
Evaluation of protein sequence encodings. One-hot encodings outperform other simple protein representations on a held-out test dataset. Neural encodings from pretrained protein language models slightly outperform one-hot encodings on a held-out test dataset. Metrics were averaged across 10 random seeds.

**Table S3.** Evaluation of the state-of-the-art model relative to ADhunter. ADhunter outperforms the stateof-the-art AD predictor, TADA, when trained on the initial dataset and evaluated on a held-out test dataset. PADDLE was not included since the untrained model is not available for fair comparison.

| Model | Pearson r | RMSE |
| --- | --- | --- |
| TADA | 0.538 | 0.995 |
| ADhunter | 0.775 | 0.631 |

**Table S4.** TADA ablation study. We evaluated the contribution of each architectural component of TADA by incrementally removing or replacing key elements then training on the initial dataset and evaluating on a held-out test dataset. Metrics were averaged across three random seeds.

| Model variant | Accuracy | F1 score |
| --- | --- | --- |
| Conv1D-Dropout-Conv1D-Dropout-Attention-BiLSTM-BiLSTM | 0.845 | 0.845 |
| Conv1D-Dropout-Conv1D-Dropout-Attention-BiLSTM | 0.845 | 0.845 |
| Conv1D-Dropout-Conv1D-Dropout-Attention-MaxPooling1D | 0.839 | 0.839 |
| Conv1D-Dropout-Conv1D-Dropout-MaxPooling1D | 0.852 | 0.852 |
| Conv1D-Dropout-MaxPooling1D | 0.849 | 0.849 |
| Conv1D-Dropout-Conv1D-Dropout-BiLSTM-BiLSTM | 0.850 | 0.850 |
| Conv1D-Conv1D-Attention-BiLSTM-BiLSTM | 0.846 | 0.846 |
| Conv1D-Conv1D-BiLSTM-BiLSTM | 0.852 | 0.852 |
| Conv1D-Conv1D-MaxPooling1D | 0.866 | 0.866 |
| Conv1D-MaxPooling1D | 0.860 | 0.860 |
| Attention-MaxPooling1D | 0.839 | 0.838 |
| BiLSTM | 0.863 | 0.863 |

**Table S5.** Evaluation of model generalizability. Using spectral clustering analysis, the ensemble model of ADhunter with neural encodings outperforms single models of ADhunter with simple (one-hot) or neural (ESM) encoding at generalizing on the initial dataset. Metrics were averaged across three random seeds.

| Model | Pearson r | RMSE |
| --- | --- | --- |
| TADA | 0.381 | 1.021 |
| ADhunter_simple | 0.416 | 0.884 |
| ADhunter_neural | 0.476 | 0.856 |
| ADhunter | 0.512 | 0.826 |

**Table S6.** ADhunter achieves state-of-the-art performance as a surrogate model for AD activity. ADhunter was compared to the state-of-the-art AD predictor, TADA, when trained and tested on either the harmonized dataset or the Morffy et al. dataset. ADhunter outperformed TADA on both datasets as well as regression and classification prediction tasks. Metrics are averaged across three random seeds.

| Prediction Task | Model | Harmonized Dataset | Morffy et al. Dataset |
| --- | --- | --- | --- |
| Regression | TADA | Pearson r = 0.621;<br>RMSE = 0.935 | Pearson r = 0.635;<br>RMSE = 0.961 |
|  | ADhunter | Pearson r = 0.818;<br>RMSE = 0.556 | Pearson r = 0.682;<br>RMSE = 0.740 |
| Classification | TADA | Accuracy = 0.861;<br>F1 score = 0.866 | Accuracy = 0.924;<br>F1 score = 0.922 |
|  | ADhunter | Accuracy = 0.905;<br>F1 score = 0.910 | Accuracy = 0.932;<br>F1 score = 0.935 |

## Supplementary Data

**Data S1.** Initial dataset.

**Data S2.** Active learning library.

**Data S3.** Harmonized dataset.

**Data S4.** Oligonucleotides used in this study.

**Data S5.** Fluorescence-activated cell sorting metrics.

## Notes

https://github.com/shih-lab/ADhunter

https://www.ncbi.nlm.nih.gov/bioproject/PRJNA1183837/

### Summary of Updates

The title has been updated; author affiliations have been updated for completeness; the font has been changed from serif to sans serif; an error where the taxonomical labels were not shown in Figure 2C has been fixed.

